# Betaproteobacterial clade II *nosZ* activated under high N_2_O concentrations in paddy soil microcosms

**DOI:** 10.1101/2025.02.17.638610

**Authors:** Kazumori Mise, Yoko Masuda, Keishi Senoo, Hideomi Itoh

## Abstract

**Aims:** Microbial communities in paddy soils act as potential sinks of nitrous oxide (N_2_O), a notorious greenhouse gas, but their potential to reduce external N_2_O is unclear. The direct observation of N_2_O reduction in submerged field soils is technically difficult. Here, we aimed to identify soil microbial clades that underpin the strong N_2_O mitigation capacity.

**Methods and Results:** We constructed paddy soil microcosms with external N_2_O amendment that enabled the simultaneous evaluation of N_2_O reductase gene (*nosZ*) transcripts and N_2_O consumption. Although the amount of N_2_O amended was large, it was mostly consumed after 6– 8 days of microcosm incubation. Metatranscriptomic sequencing revealed that betaproteobacterial *nosZ*, especially those classified as clade II *nosZ* belonging to the orders *Rhodocyclales* or *Nitrosomonadales*, occupied >50% of the *nosZ* transcripts in three of the five paddy soils used. On the other hand, publicly available shotgun metagenomic sequences of 46 paddy soils were not dominated by betaproteobacterial clade II *nosZ* sequences, although they were ubiquitous. The same applied to the 16S rRNA sequences of *Rhodocyclales* or *Nitrosomonadales*.

**Conclusions:** The results indicated that betaproteobacterial N_2_O reducers potentially serve as powerful N_2_O sinks. *Betaproteobacteria* holding clade II *nosZ* can be targets of biostimulation, although further studies are required to understand their ecophysiology.

**Impact Statement:** Our results indicate that *Rhodocyclales* or *Nitrosomonadales* have the potential to serve as powerful N_2_O sinks in soils.

## INTRODUCTION

Nitrous oxide (N_2_O) is a potent greenhouse gas, with a global warming potential approximately 265 times that of carbon dioxide (Pachauri et al., 2014). N_2_O depletes ozone and is expected to become the dominant ozone-depleting compound of this century, surpassing chlorofluorocarbons and hydrochlorofluorocarbons, for which countermeasures have been developed (Ravishankara et al., 2009). Its long atmospheric lifetime, typically spanning over a century, allows it to persist in the atmosphere and exert a lasting impact on both global warming and ozone depletion. Over the past four decades, anthropogenic N_2_O emissions have increased by >40%, primarily driven by agricultural activities, especially the application of nitrogen fertilizers (Tian et al., 2023). Currently, >30% of anthropogenic N_2_O emissions are ascribed to agricultural soils (Tian et al., 2023), a proportion that is expected to increase with continued chemical fertilizer use, which triggers microbial processes, such as nitrification and denitrification that release N_2_O (Shcherbak et al., 2014). Natural soils contribute more than one-half of global N_2_O emission from nature (Tian et al., 2023). In short, soil is a major source of N_2_O emissions.

Flooded paddy soils are agricultural soils with relatively low N_2_O emissions (Bouwman et al., 2002; Yan et al., 2003; Nishimura et al., 2005; Akiyama et al., 2006). Under anoxic conditions, which develop in flooded paddy soils, electron acceptors other than oxygen play vital roles, resulting in high denitrification activity, which includes both N_2_O production and consumption (NO_3_⁻ → NO_2_⁻ → NO → N_2_O → N_2_) (Kuypers et al., 2018). Indeed, >97% of the N_2_O produced in paddy soil is consumed without release to the atmosphere (Wang et al., 2022), and this tendency has been observed regardless of the soil type (Wang et al., 2024a). These suggest that paddy soils exhibit high N_2_O reduction activity. Thus, paddy soils are not only characterized by low N_2_O emissions but also increasingly recognized for their ecological function as N_2_O sinks (Zhong et al., 2023; Wang et al., 2024b). However, which microbes underpin this strong N_2_O mitigation capacity in paddy soil is unclear.

N_2_O-reducing microbes comprise phylogenetically diverse clades of bacteria, and the kinetics of N_2_O-reducing activity is also diverse (Hiis et al., 2024). Microbial N_2_O reduction is driven by N_2_O-reductase, NosZ, which is encoded by *nosZ* gene (Kuypers et al., 2018). Currently known *nosZ* sequences are classified into two distinct groups with different molecular machineries and phylogeny, namely clade I (also known as typical *nosZ*, *nosZI,* or TAT-dependent type) and clade II (atypical *nosZ*, *nosZII,* or Sec-dependent type). Clade I *nosZ* is distributed across *Alphaproteobacteria*, *Betaproteobacteria*, and *Gammaproteobacteria*, whereas clade II *nosZ* is distributed among more diverse lineages of bacteria, including *Bacteroidota*, *Gemmatimonadota*, *Myxococcota*, *Verrucomicrobiota*, and some clades of *Betaproteobacteria* (Jones et al., 2013). Metagenomic studies have indicated that soils harbor diverse lineages of *nosZ* holders, with a dominance of clade II *nosZ* (Orellana et al., 2014; Nadeau et al., 2019). A metatranscriptomic study using field soils reported that *Myxococcota nosZ* (presumably belonging to clade II) is highly abundant in waterlogged paddy soils (Masuda et al., 2017). In addition to soils, activated sludge and wastewater treatment plants are dominated by clade II *nosZ* transcripts (Li et al., 2021).

Although many studies have reported community structures of N_2_O reducers in paddy soils, their potential to reduce external N_2_O is unclear. Direct observation of N_2_O dynamics in paddy fields is technically challenging under flooded conditions (Masuda et al., 2019). The only feasible method to simultaneously observe N_2_O reduction and N_2_O-reducing community structures would be to use closed systems, such as soil microcosms.

Previous studies have utilized microcosm experiments with labile carbon sources and DNA/PCR-based analyses to investigate the microbial communities responsible for N_2_O reduction in paddy soils (Ishii et al., 2011; Qin et al., 2020; Xing et al., 2021; Maheshwari et al., 2023; Shaaban et al., 2023; Xiang et al., 2023). However, these DNA-based approaches primarily reflect microbial potential rather than actual activity, because they do not capture gene expression or microbial functions in real-time (Prosser, 2015). PCR-based methods are susceptible to biases, including the preferential amplification of certain sequences and variations in primer specificity and amplification efficiency (Polz and Cavanaugh, 1998; Ruijter et al., 2009; Jones et al., 2013; Masuda et al., 2024). The external input of labile carbon sources, such as glucose or succinate, may disturb microbial communities in a way that is unrealistic in field soils. To address these limitations, using carbon sources more representative of the natural environment or RNA-based analyses can offer a more sensitive and accurate method to identify microorganisms that actively respond to N_2_O. RNA-based approaches, such as metatranscriptomics, provide insights into the gene expression of N_2_O reducers, offering a better understanding of their role in N_2_O reduction and improving the overall assessment of microbial activity in paddy soils.

Here, we aimed to identify paddy soil microbes that can reduce a substantial amount of external N_2_O. We constructed soil microcosms amended with rice straw and N_2_O and obtained time-series metatranscriptomic profiles while monitoring the amount of N_2_O consumed, to identify active members within N_2_O-reducing communities. To put the results into a more general context, we analyzed the metatranscriptome of N_2_O-reducing communities constructed using paddy soils from four distinct locations. We further investigated the global distributions of these actively expressed *nosZ* through a meta-analysis of soil metagenomic datasets collected worldwide.

## EXPERIMENTAL PROCEDURES

### Soil sampling

The soils used in this study, named Soils X or Y1–Y4, were taken from the surfaces of five paddy fields in Japan (Table 1). Soil X was from a long-term fertilization field managed by a public research organization, with its physicochemical and microbiome profiles intensively investigated (Itoh et al., 2013; Masuda et al., 2023). Soils Y1–Y4 were provided by private farmers. The sampled soils were stored at 4 °C until use. The physicochemical properties of soil were determined, as described previously (Masuda et al., 2024). In brief, soil pH(H_2_O) and electrical conductivity were determined in the water suspension (soil:water = 1:5 [w/w]), whereas soil total carbon and nitrogen were measured using the dry combustion method.

**Table 1.**
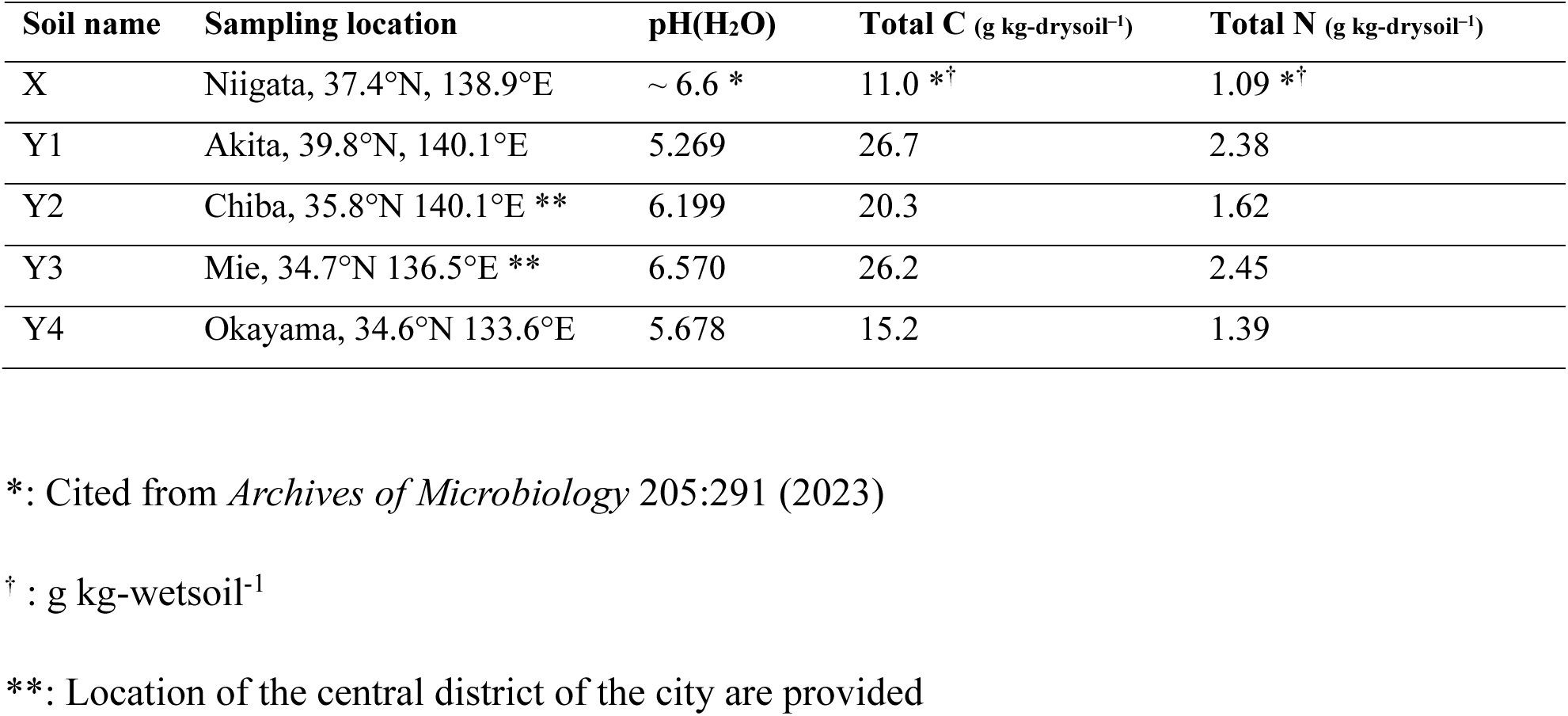
Descriptions of soil samples used in this study.

| Soil name | Sampling location | pH(H<sub>2</sub>O) | Total C (g kg-drysoil<sup>-1</sup>) | Total N (g kg-drysoil<sup>-1</sup>) |
| --- | --- | --- | --- | --- |
| X | Niigata, 37.4°N, 138.9°E | ~ 6.6 * | 11.0 *<sup>†</sup> | 1.09 *<sup>†</sup> |
| Y1 | Akita, 39.8°N, 140.1°E | 5.269 | 26.7 | 2.38 |
| Y2 | Chiba, 35.8°N 140.1°E ** | 6.199 | 20.3 | 1.62 |
| Y3 | Mie, 34.7°N 136.5°E ** | 6.570 | 26.2 | 2.45 |
| Y4 | Okayama, 34.6°N 133.6°E | 5.678 | 15.2 | 1.39 |

\*: Cited from *Archives of Microbiology* 205:291 (2023)
<sup>†</sup> : g kg-wetsoil<sup>-1</sup>
\*\* : Location of the central district of the city are provided

### Soil microcosm incubation

We first constructed dozens of microcosms using Soil X (Table 1) and 10-mL glass vials (with approximately 13 mL of inner capacity). We mixed the wet soils and distilled water and preincubated the sieved mixture at 25 °C without light for 7 days. Soils were sieved through a 2-mm mesh to remove large plant residues and gravel. Then, we adjusted the water content so that each microcosm consisted of 2.5 g of wet soil (water content = 40% [w/w]) and 1.0 mL of distilled water. This means that the final water content was (2.5 × 40% + 1.0)/(2.5 + 1.0) = 57.1%. Each microcosm was amended with 25 mg of dried rice straw powder (<0.25 mm in length) as possible electron donors (Eusufzai et al., 2011; Zhang et al., 2023). The microcosm vials were tightly sealed with butyl rubber stoppers and aluminum caps, with their headspaces completely replaced with N_2_O and argon (Ar) gas. We applied N_2_O and Ar to some of the microcosms with N_2_O concentrations of 4.46 mM, whereas the others were amended only with Ar. We defined this point as Day 0. Glass vials with 3.5 mL of sterilized distilled water, with N_2_O-Ar gas (N_2_O concentration: 4.46 mM) amended in their headspace gas phases, were prepared as negative controls.

After 0–7 days of incubation at 25 °C without light, the microcosm soils were destructively sampled for measuring headspace N_2_O content or for metatranscriptomic sequencing. Four N_2_O-added microcosms were sampled for metatranscriptomics every day, and three microcosms without added N_2_O were sampled on Days 2, 4, and 7. For N_2_O measurement, four microcosms with and without N_2_O addition (eight in total) were destructively sampled and analyzed using a gas chromatograph, as described previously (Kuroda et al., 2022).

For comparison, we constructed three microcosms using each of Soils Y1–Y4 (Table 1). The microcosms were prepared in the same manner as those for Soil X, with the final water content adjusted to ∼57.1% (the same as Soil X). In total, 20 mg of straw was applied. All 12 microcosms received N_2_O amendment, and the N_2_O content of each microcosm was monitored using a GC3210 Gas Chromatograph (GL Science, Tokyo, Japan) and a column packed with ShinCarbon-ST 50/80 (Shinwa Chemical Industries, Kyoto, Japan). The temperatures of the TCD and columns were set at 210 and 150 °C, respectively. The flow rate of the carrier gas was 30 mL/min. The microcosms were destructively sampled for metatranscriptomic sequencing when they largely ran out of spiked N_2_O. The microcosms of Soils Y1, Y2, Y3, and Y4 were sampled on Days 8, 7, 6, and 8, respectively.

### RNA extraction and metatranscriptomic sequencing

RNA was extracted from each microcosm immediately after sampling using RNesay PowerSoil Total RNA Kit (Qiagen), according to the manufacturer’s protocol. Concomitant DNA was eliminated using TURBO DNA-free Kit (ThermoFisher). The resulting RNA solutions were purified and concentrated using RNA Clean & Concentrator-5 (ZYMO RESEARCH). Bacterial rRNA was eliminated using riboPool (siTOOLs Biotech), followed by sequencing library construction using MGIEasy RNA Directional Library Prep Set (MGI Tech) with a fragmentation step at 82 °C for 8 minutes. The quantity and quality of the constructed libraries were verified using Synergy H1 (Bio Tek), QuanitFluor dsDNA System (Promega), Fragment Analyzer (Advanced Analytical Technologies), and Bioanalyzer (Agilent Technologies). The libraries were transformed into DNA nanoballs (DNBs) using MGIEasy Circularization Kit (MGI Tech) and DNBSEQ-G400RS High-throughput Sequencing Kit (MGI Tech). DNBs were sequenced using DNBSEQ-G400 in the 200-bp paired-end mode.

### Bioinformatic analyses of metatranscriptomic reads

Metatranscriptomic reads underwent the following analyses. Adaptor sequences were trimmed off each read using Cutadapt v4.8 (Martin, 2011) with default parameter settings. The low-quality region of each read was eliminated using vsearch v2.15.2_linux_x86_64 (Rognes et al., 2016) with the options “--fastq_truncee 1 --fastq_minlen 50”. We eliminated reads bearing rRNA gene sequences using SortMeRNA v4.3.6 (Kopylova et al., 2012) with the default reference database and “--paired_in” option. The filtered reads were finally assembled using SPAdes v3.15.5 (default parameters under “--rna” flag) (Bushmanova et al., 2019; Prjibelski et al., 2020). Reads obtained from the same soil (listed in Table 1) were assembled together (i.e., coassembled), resulting in five sets of contigs. Regarding Soil X, which yielded 41 samples and >4.7 × 10^11^ bases of metatranscriptomic reads, we randomly selected 5,000,000 read pairs per sample (quality-filtered reads only) and used them for assembly to save computational resources. Note that this subsampling applies only to the assembly process; the read depth of each contig or coding sequence (CDS) was calculated using all reads (details explained later in this section).

CDS predictions and functional annotations were given to contigs of ≥500 bp, using eggNOG-mapper v2.1.2 (Cantalapiedra et al., 2021) and the default reference database (Huerta-Cepas et al., 2019), with the parameter settings of “--itype proteins --tax_scope_mode inner_narrowest --tax_scope 1 --target_taxa 1”. Contigs of <500 bp were discarded and not used for downstream analyses. For CDSs predicted to encode NosZ (K00376 in KEGG orthology, (Kanehisa et al., 2024)), we performed a more stringent phylogenetic and taxonomic annotation using phylogenetic placement implemented in pplacer v1.1.alpha19-0-g807f6f3 (Matsen et al., 2010) and a custom database of NosZ sequences. We also used MAFFT v7.525 (Katoh et al., 2002) for phylogenetic placement. The construction of the custom NosZ database was dependent on Prodigal (Hyatt et al., 2010), FunGene (Fish et al., 2013), KOfam (Aramaki et al., 2020), HMMER (Eddy and Wheeler, 2007), and GTDB (Parks et al., 2022). The full details are explained in Supplementary Text 1.

To evaluate the abundance of the transcripts of each gene, we mapped all the preassembled reads back onto the contigs using BWA-MEM2 version 2.2.1 (Vasimuddin et al., 2019). We then calculated transcripts per kilobase million (TPM) of each CDS using “bedcov” command implemented in SAMtools version 1.18 (Danecek et al., 2021). The CDSs annotated as betaproteobacterial *nosZ*, with a TPM of ≥ 5.0 in at least one sample, were aligned using MAFFT, and their phylogenetic tree was constructed using FastTree version 2.1.11 with “pseudo” option (Price et al., 2009). We also performed canonical bootstrapping test (100 times) using a custom script for randomizing alignments (see Data Availability) and FastTree.

### DNA extraction and metagenomic analyses

We obtained shotgun metagenomic sequences of the soils used in the microcosm experiments. Metagenomic sequencing of Soil X has been reported in a previous study—“SF” samples described in Masuda et al. (2023)—therefore, we sequenced the metagenomes of Soils Y1–Y4. DNA was extracted from approximately 0.25 g of each soil using DNeasy PowerSoil Pro Kit (Qiagen). The extracted DNA samples were sequenced on DNBSEQ following the protocol established by the manufacturer with slight modifications (Masuda et al., 2024). The metagenomic reads underwent adaptor trimming and quality filtering as described previously (Masuda et al., 2024). Regarding the metagenomic data of Soil X, all reads from the three biological replications were pooled and treated as one sample in this study.

### Global distribution and diversity of *nosZ* sequences in metagenomes

In addition to the metagenomic sequence data described above, we used intermediate data obtained in a meta-analysis study (https://plus.figshare.com/articles/dataset/Quality-filtered_soil_shotgun_metagenomes/25332547 from Masuda et al., 2024). This dataset includes quality-filtered sequences from 41 shotgun metagenomic datasets from paddy soils collected worldwide, in addition to other metagenomic sequences. We analyzed the metagenomic data to investigate the distribution and ubiquity of betaproteobacterial *nosZ* sequences, which were abundant in our metatranscriptomic sequences (see Results).

To search for *nosZ* from the metagenomic reads with minimal computational costs, we performed a two-step homology search. First, all reads were subjected to homology search against the NosZ database constructed from the eggNOG database. This database consists of all protein sequences with annotations of K00376 in the eggNOG database and is compatible with a large number of query sequences because of its small size. Here we used the blastx command in DIAMOND v2.1.9.163, with the options “-e 1e-5 -k 1”. Query sequences that had a significant hit were regarded as candidates for *nosZ* sequences. To select “true” *nosZ* sequences, they were mapped onto the whole size of the eggNOG database using the blastx command in DIAMOND (options: -e 1e-5 -k 200). Although we used only the top hits to identify *nosZ* sequences, we obtained the top 200 hits per query to mitigate the inaccuracy resulting from the heuristics of DIAMOND. The determined *nosZ* reads were further subjected to a more stringent phylogenetic annotation. See Supplementary Text 1 for details.

Furthermore, we calculated the proportion of 16S rRNA sequences belonging to order *Rhodocyclales* or *Nitrosomonadales* in each metagenome. The 16S rRNA sequences were picked and annotated using SortMeRNA, blastn search, SINTAX (Edgar, 2016), and SILVA v138.1, as described previously (Masuda et al., 2024). Part of this analysis has been reported in a previous work (Masuda et al., 2024). We reused the intermediate results that are available in FigShare (https://figshare.com/articles/dataset/Microbiome_12_95/25894636).

Throughout the study, we used SeqKit (Shen et al., 2016), TaxonKit (Shen and Ren, 2021), The Newick Utilities (Junier and Zdobnov, 2010), and R v4.0.5 (R Core Team https://www.R-project.org/) to handle fastq/fasta files, to manage NCBI taxonomy IDs, to edit newick files, and to visualize data, respectively.

## RESULTS

### N_2_O consumption in Soil X

We first overview the time-series datasets obtained from Soil X microcosms. The temporal changes in N_2_O concentration in the microcosms are shown in Fig. 1a. N_2_O concentration at Day 0 was 3.53 ± 0.15 mM (mean ± SD), which is 21% lower than the due concentrations of amended N_2_O. The amount of lost N_2_O was 9.3 ± 1.5 μmol, which is 8.8 times lower than the solubility of N_2_O in water (1.2 g/L = 82 μmol per 3 mL of water). Despite the high concentration of N_2_O applied, inoculated N_2_O was largely consumed in the fresh-soil microcosms within 7 days of incubation. The rate of N_2_O consumption gradually increased with a peak at Days 2–4, and then decreased until N_2_O ran out (Fig. 1a). Although N_2_O is highly soluble in water, such a decline in N_2_O concentration was not observed in microcosms consisting only of water without soil (Fig. 1a).

**Figure 1.**
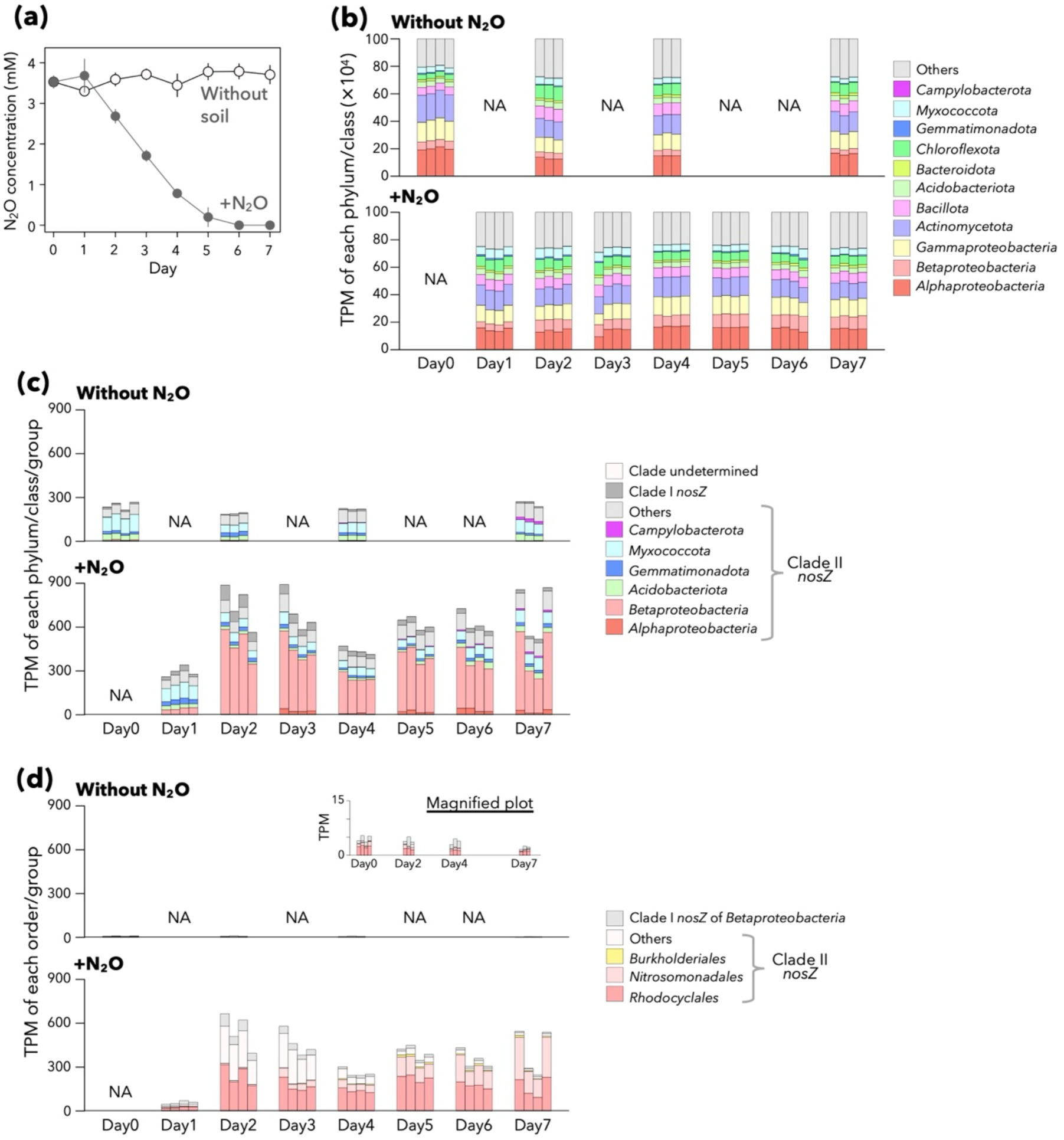
Overview of the results of the microcosm experiments using Soil. **X.** (a) Dynamics of N_2_O concentrations in the microcosm headspaces. Results for samples with and without N_2_O amendment are displayed in closed and open circles, respectively. The average of four biological replications is indicated. Error bars indicate standard deviations. Some of the error bars are shorter than the circles. (b) Dynamics of the overall taxonomic composition of the metatranscriptomic sequences. All bacterial contigs that received valid annotations were considered. The transcripts per kilobase million (TPMs) of bacterial phyla and proteobacterial classes are indicated. Each bar indicates data from one microcosm (i.e., one biological replication). NA denotes not available. (c) Dynamics of the *nosZ* TPM in the metatranscriptomic sequences. The scales are the same between the two panels, each representing samples without and with N_2_O amendment. NA denotes not available. (d) Dynamics of betaproteobacterial *nosZ* TPM in the metatranscriptomic sequences. The scales are the same between the two panels, although a magnified plot of the upper panel is presented. NA denotes not available.

### Time-series transitions of metatranscriptomic profiles

A total of 1,181,867,200 read pairs (22,549,292–33,206,316 per sample) were obtained in high-throughput sequencing, and 1,001,865,287 (84.8%) passed quality filtering and rRNA removal. *De novo* assembly of the filtered sequences yielded 3,699,693 contigs, and 1,002,468 (27.1%) were ≥500 bp (the remainders were discarded in downstream analysis). Functional and phylogenetic annotations using eggNOG-mapper indicated that 854,624 (85.3%) encoded at least one prokaryotic gene, and 483 encoded *nosZ* (484 *nosZ* CDSs in total). Phylogenetic analysis indicated that 438 (90.7%) of *nosZ* belonged to clade II (also known as “atypical” *nosZ*) rather than clade I (“typical” *nosZ*).

Although most of the *nosZ*-harboring contigs were <2000 bp in length and their operon structures were not observed, longer contigs presented conserved structures of *nos* operons. In particular, several long contigs bearing *Rhodocyclales* clade II *nosZ* were obtained. They presented conserved structures, with *cytC1*, *nosD*, *nosH*, and *nosF* encoded immediately downstream (some examples are shown in Fig. S1). They are congruent with the known structures of *nos* operons in *Rhodocyclales* genomes (Sanford et al., 2012; Semedo et al., 2020; He et al., 2024). The same applies to the *nosZ* of *Nitrosomonadales*, a group phylogenetically close to *Rhodocyclales*. For comparison, clade I *nosZ* was adjacent to *nosRDFYL*., which is also consistent to known structures of clade I *nosZ* (Sanford et al., 2012).

We mapped the metatranscriptomic reads onto the contigs, and a large portion of the metatranscriptomic reads (78.4%–94.2%) were successfully mapped. The time-series metatranscriptomic profiles showed that the taxonomic compositions of the transcripts were overall similar during the 7-day incubation (Fig. 1b), whereas the expression of *nosZ* was dynamically promoted in samples with N_2_O amendment (Fig. 1c). The TPM of *nosZ* increased by 3.05 times on Day 2 with N_2_O applied (compared with Day 0), whereas it remained stable in microcosms without N_2_O amendment. This result aligns with the contemporary decrease in N_2_O concentration: the expression of *nosZ* and the decrease in N_2_O concentration coincided. At this moment, we observed increased expression of accessory genes that are encoded in the neighbors of clade II, such as *nosD*, *nosH*, and *nosF* (Fig. S2).

### Betaproteobacterial *nosZ* dominated the metatranscriptome of Soil X

The phylogeny of expressed *nosZ* spanned diverse bacterial clades, with the dominant phyla of *Pseudomonadota*, *Acidobacteriota*, and *Myxococcota* encoding clade II *nosZ* (Fig. 1c). By contrast, the increase in overall *nosZ* expression was largely attributed to *Betaproteobacteria*, with dominance of order *Rhodocyclales*. The TPM of clade II *nosZ* annotated as *Betaproteobacteria* increased by 69.8 times between Days 0 and 2, respectively. The same applied to the TPM of *Rhodocyclales nosZ* (Fig. 1d). At the later stages of incubation, particularly after Day 5, the TPM of *Nitrosomonadales* clade II *nosZ* increased. *Nitrosomonadales* (also known as *Spirillales* in ICNP nomenclature) belongs to *Betaproteobacteria* and is phylogenetically closer to *Rhodocyclales* than other subgroups (Boden et al., 2017). Regarding lineages other than *Betaproteobacteria*, N_2_O amendment did not have a major impact on their expression levels. In addition, those of other *nosZ* did not present such drastic changes (Fig. 1c).

Most of the *nosZ* of *Rhodocyclales* belonged to clade II *nosZ*, whereas clade I *nosZ* of *Rhodocyclales* was encoded on only two contigs. By contrast, the *nosZ* sequences of *Rhodocyclales* or *Nitrosomonadales* were not phylogenetically uniform and comprised multiple genera or species. Several clades of *nosZ* presented two distinct temporal patterns: some peaked at Day 2 and gradually decreased by Day 4, whereas others presented a major increase at a later stage (Fig. 2). These two types of *nosZ* were phylogenetically conserved, although it should be noted that the phylogenetic tree presented low bootstrap values (Fig. 2). Overall, although N_2_O-reducing traits are conserved within *Rhodocyclales* and *Nitrosomonadales*, they exhibit some diversity.

**Figure 2.**
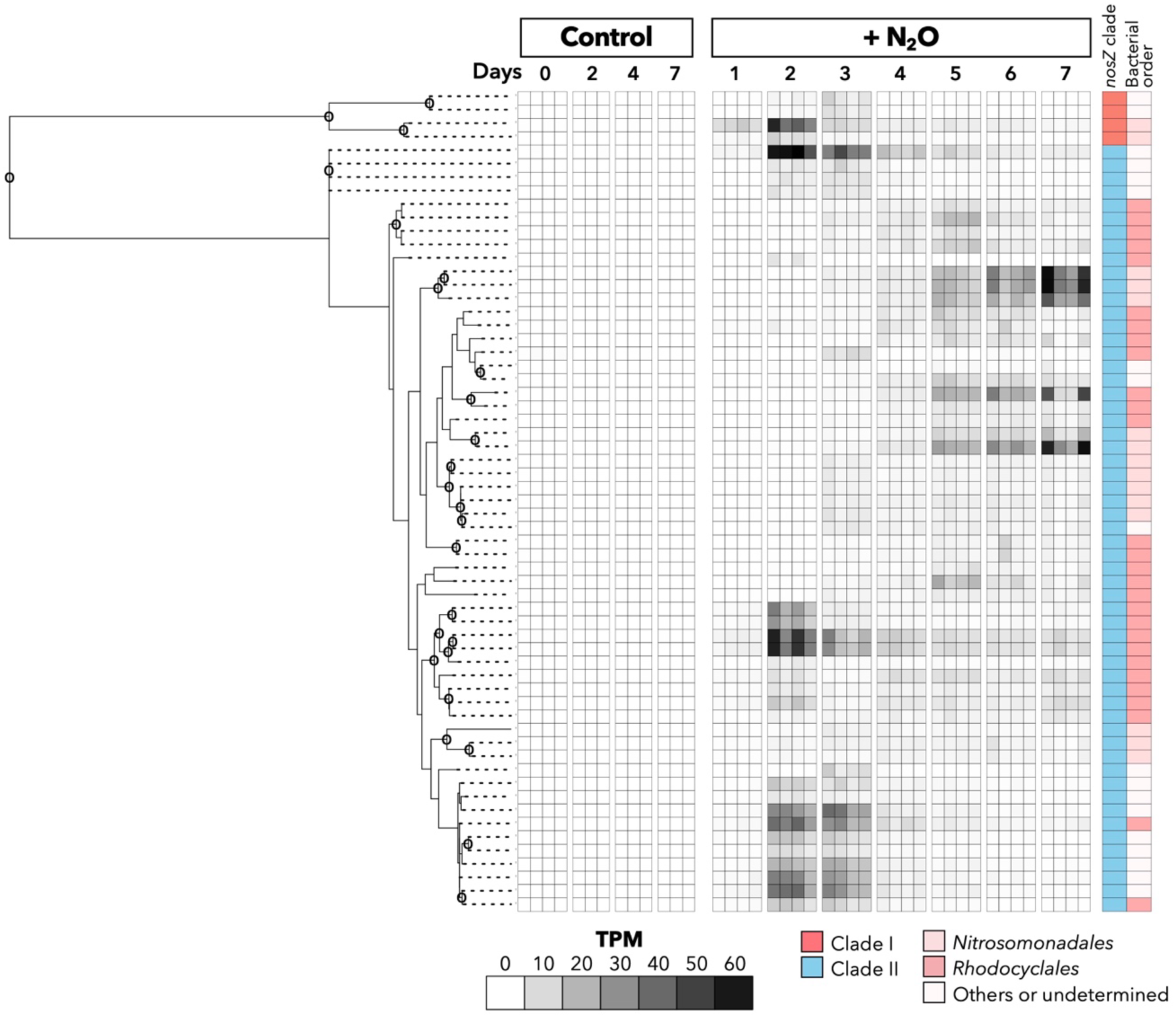
Phylogeny and detailed dynamics of betaproteobacterial *nosZ* expression in Soil X microcosms. The tree dendrogram represents an approximate maximum-likelihood phylogeny of betaproteobacterial NosZ encoded on metatranscriptomic contigs. Circles indicate canonical bootstrap values of 70% or higher. Note that the bootstrap values are different from “local support values” implemented in FastTree. The heatmap indicates the transcripts per kilobase million (TPM) of each CDS. The two rightmost stripes represent the clade and taxonomic classification of each NosZ. Only betaproteobacterial *nosZ* CDSs with a TPM of ≥ 5.0 in at least one sample are shown.

The TPM of all *Rhodocyclales* transcripts (including but not limited to *nosZ*) steadily increased, indicating that the increased transcription of *Rhodocyclales nosZ* supported their growth and the inoculated N_2_O served as the terminal electron acceptor. By contrast, the overall expression profiles (summarized by KEGG pathways) were not extensively affected by N_2_O inoculation (Fig. 3). They depended more on different incubation periods than whether N_2_O was inoculated.

**Figure 3.**
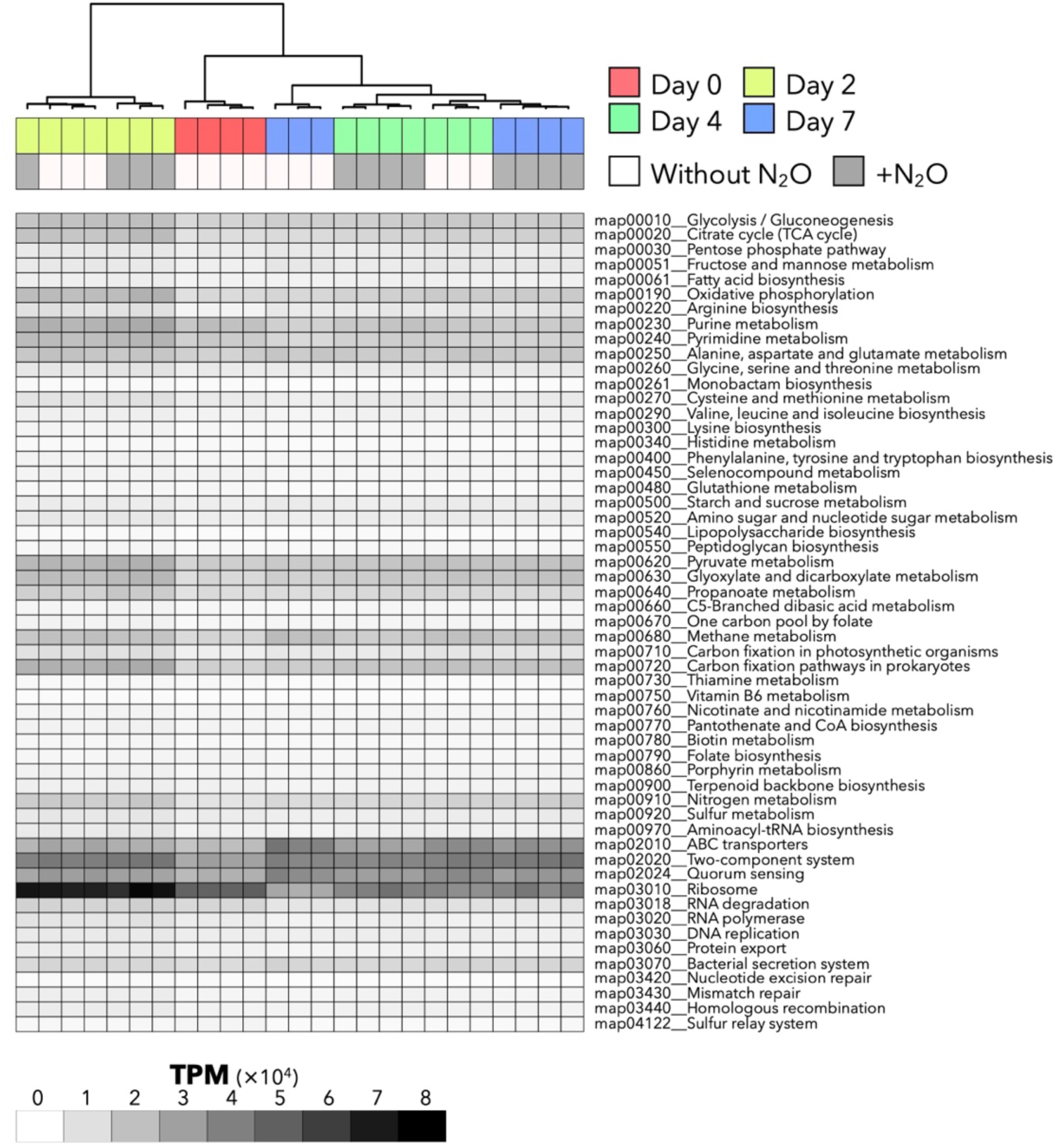
Comparison of the overall metatranscriptomic profiles among the Soil X microcosms. The heatmap indicates the TPM of each pathway (KEGG pathways). Each column represents one sample, and the sample attributes are indicated above the heatmap. Clustering was performed using Ward’s method with Manhattan distances.

### Atypical *nosZ* and *Rhodocyclales* dominate in microcosms of different paddy soils

To investigate if the observations in Soil X are applicable to other paddy soils, we performed similar microcosm experiments using four types of paddy soils (Soils Y1–Y4; Table 1). In all soils, N_2_O was rapidly consumed within a week (Fig. 4a). At the time when microcosms largely ran out of N_2_O, metatranscriptomic analyses proved the transcription of diverse lineages of *nosZ*. TPMs of *nosZ* in the four soils were comparable to those of Soil X (350.0–1260.4, Fig. 4b). Other statistical summaries of metatranscriptomic sequencing and assemblies can be found in Table S2.

**Figure 4.**
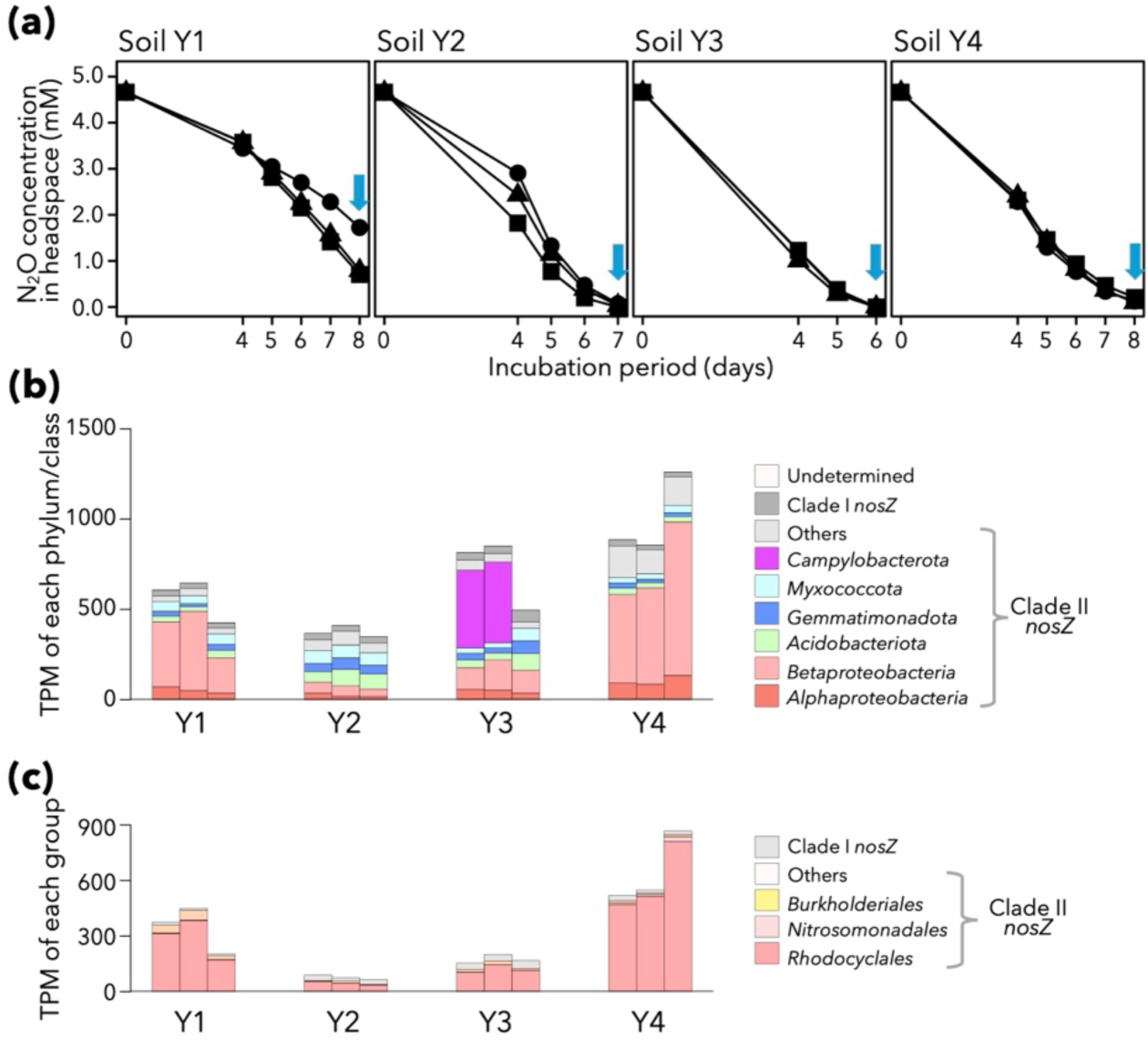
Overview of the results of the microcosm experiments using Soils Y1–Y4. (a) Dynamics of N_2_O concentrations in the microcosm headspaces. Blue arrows indicate the time point for metatranscriptomic sequencing. (b) Abundance and diversity of *nosZ* reads in the metatranscriptomic sequences. (c) Abundance and diversity of betaproteobacterial *nosZ* reads in the metatranscriptomic sequences.

Most of the *nosZ* sequences belonged to clade II, particularly those of *Betaproteobacteria* and *Rhodocyclales*, with the exception of Soil Y2. This trend was similar to Soil X microcosms. By contrast, the expression of *Nitrosomonadales nosZ* was relatively minor in all four soils. The other dominant *nosZ* holders included *Alphaproteobacteria*, *Acidobacteriota*, *Gemmatimonadota*, and *Myxococcota*. *Campylobacterota nosZ* were dominant in two of the microcosms with Soil Y3, but were not observed in other soils.

### *nosZ*-holding *Betaproteobacteria* are not dominant in soil microbiomes

We asked whether betaproteobacterial *nosZ* are prevalent and dominant in soils in nature, especially in paddy soils. We performed a semiglobal-scale meta-analysis of 46 paddy soil samples (mostly from Asia), which comprises 41 samples analyzed in Masuda et al. (2024) and the five raw soils used for our microcosm experiments. The geographic origins of the 46 samples are displayed in Fig. 5a. Each sample yielded 72–20,704 reads of *nosZ* and 1,023–253,450 reads of 16S rRNA genes.

**Figure 5.**
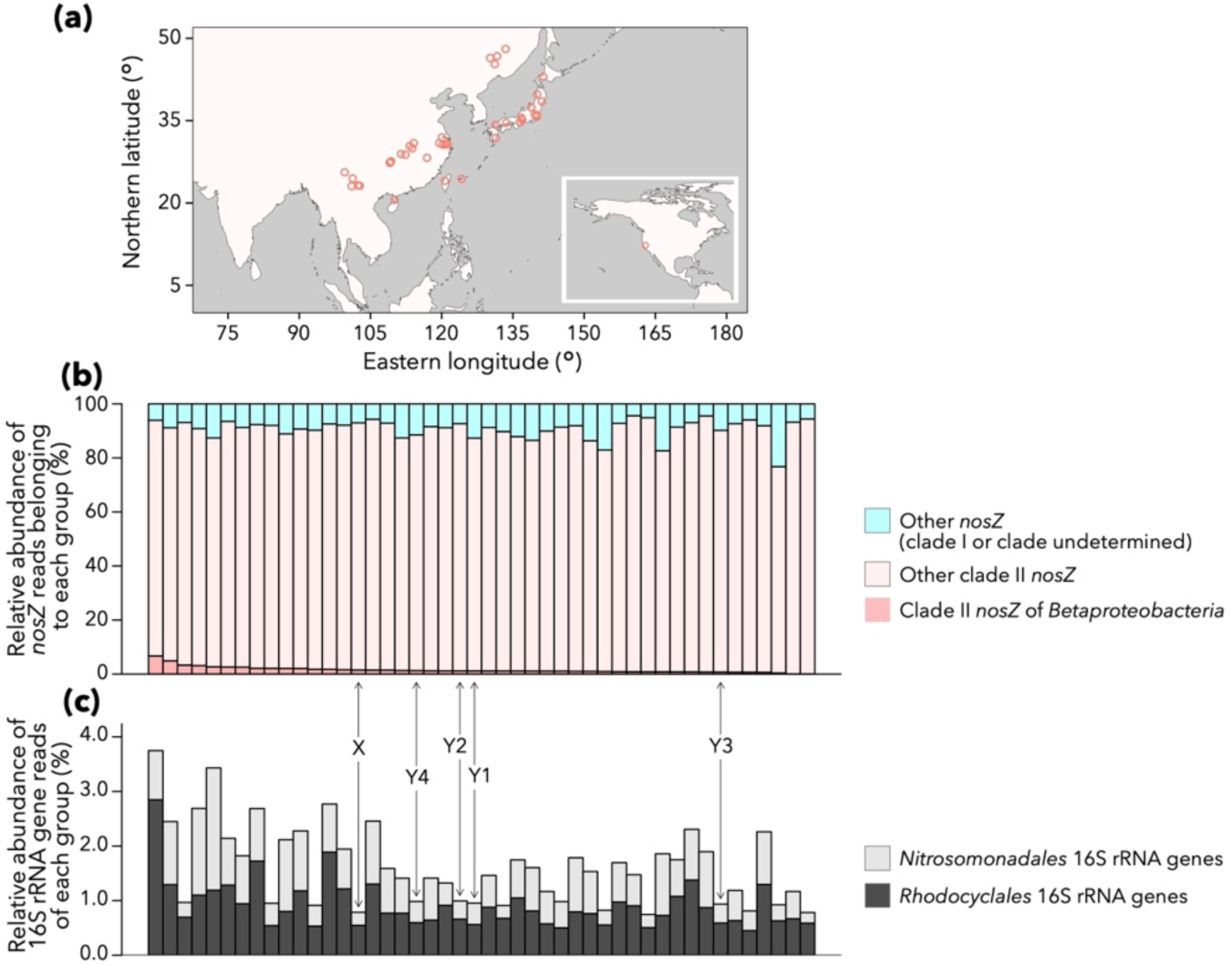
Distribution of betaproteobacterial *nosZ* in soil metagenomic sequences collected worldwide. (a) Geographic origins of the paddy soil metagenomic data used in this study. (b) Relative abundances of *nosZ* reads belonging to clade II and/or *Betaproteobacteria*. Each bar indicates the composition of one sample. Bars are sorted by the proportion of clade II betaproteobacterial *nosZ*. Arrows indicate the results for the five soils used in our microcosm experiments. (c) Relative abundances of 16S rRNA gene reads annotated as *Rhodocyclales* or *Nitrosomonadales*. The bars are sorted in the same order as that in panel (b). Arrows indicate the results for the five soils used in our microcosm experiments.

We found that betaproteobacterial *nosZ* were not dominant in paddy soils (Fig. 5b), with some variations between samples (Fig. 5b). Clade II *nosZ* was overall dominant, in congruence with our metatranscriptomic results and previous metagenomic studies (Orellana et al., 2014). In addition, the 16S rRNA gene reads of *Rhodocyclales* and *Nitrosomonadales* were distributed ubiquitously, with the relative abundances of 0.45–2.85% and 0.20–2.24%, respectively (Fig. 5c).

## DISCUSSION

Our findings revealed that clade II *nosZ*, especially those harbored by members of *Rhodocyclales* or *Nitrosomonadales* within the class *Betaproteobacteria*, exhibited substantial transcriptional activity in N_2_O-amended soils, suggesting a key role for these taxa in reducing external N_2_O. The phylogeny of NosZ sequences (Fig. 2) and the operon structures of *nosZ*-bearing contigs (Fig. S1) indicated that the sequences are likely *bona fide nosZ*. Clade II *nosZ* was dominant in all five soils, whereas *Rhodocyclales nosZ* transcripts were dominant in four of the five soils investigated (Figs. 1d and 4bc). These trends are likely not specific to the soils used in this study, given that the five or four soils were different in geographic origins and physicochemical properties, such as pH and carbon content (Table 1). In all soils, inoculated N_2_O was consumed by Days 6–8 (Figs. 1a and 4a), although their initial concentrations being 10^5^ times higher than atmospheric levels. This rapid consumption underscores the pivotal role of clade II *nosZ*-bearing *Betaproteobacteria*, particularly *Rhodocyclales*, in N_2_O mitigation. These results align with the potential of paddy soils as N_2_O sinks, likely driven by the N_2_O-reducing capabilities of *Rhodocyclales*.

Although the prevalence of clade II *nosZ* is somewhat expected from previous metagenomic studies (Orellana et al., 2014), the dominance of *Rhodocyclales nosZ* transcripts raises several questions. First, this trend is inconsistent with the metagenomic data. Our metagenomic analysis indicated that clade II *nosZ* were consistently dominant but betaproteobacterial *nosZ*, especially those encoded by *Rhodocyclales* or *Nitrosomonadales*, were minor (Fig. 5b). In a metatranscriptomic study of soils in paddy fields, a marginal amount of betaproteobacterial *nosZ* was detected (Masuda et al., 2017). These results indicate that a limited number of *nosZ*-harboring microbes were activated in our microcosm experiments.

Second, the extreme dominance of *Rhodocyclales* clade II *nosZ* may be counterintuitive in terms of the phylogeny of *nosZ*-holding bacteria. Both clade I *nosZ* and clade II *nosZ* are distributed among *Betaproteobacteria* (including *Rhodocyclales*), and phylogenetically close members of *Rhodocyclales* can harbor different types of *nosZ*. As an extreme example, *Thauera linaloolentis* 47Lol^T^, a member of *Rhodocyclales*, harbors both types of *nosZ* in its genome (Semedo et al., 2020). Despite this, the transcripts of clade I *nosZ* from *Rhodocyclales* were not dominant, and clade II *nosZ* was selectively favored in terms of expression. Moreover, the high concentrations of amended N_2_O are unlikely to favor clade II NosZ over clade I NosZ. Clade II NosZ tends to exhibit lower *V_max_* and *K_m_* values than clade I NosZ (Yoon et al., 2016; Suenaga et al., 2019), suggesting that a high concentration of N_2_O would selectively favor bacteria holding clade I *nosZ* rather than clade II *nosZ*. This expectation, however, disagrees with our results.

Third, clade II *nosZ* are prevalent among various lineages of bacteria, including *Myxococcota* and *Acidobacteriota*. Although their 16S rRNA genes are ubiquitous and often dominant in paddy soils (Masuda et al., 2024), their expression was not promoted by N_2_O inoculation (Figs. 1b and 4b). Why only *Rhodocyclales* was selectively activated in *nosZ* holders is unclear. Although there are several factors specific to our experimental setup, they do not explain the increased expression of *Rhodocyclales nosZ*. First, high concentrations of N_2_O can have toxic effects on microbes, for example by inhibiting cobalamin activities (Sullivan et al., 2013). However, the effect of N_2_O amendment was marginal on expression profiles compared with the timepoint of sampling (Fig. 3). Second, members of *Rhodocyclales* can catabolize recalcitrant carbon substrates (Salinero et al., 2009; Xu et al., 2021); therefore, they are likely to be favored by rice straw amendment. In fact, previous studies have reported the enrichment of *Rhodocyclales* in soils amended with hydroxyethylcellulose or cellulose-rich materials (Si et al., 2018; Maheshwari et al., 2023). Similar results were observed in woodchip bioreactors, which are rich in lignocellulose (Schiml et al., 2024). In this regard, our results contrast with a soil microcosmic study using succinate as the electron donor: members of *Burlholderiales* rather than *Rhodocyclales* were presumably activated under N_2_O-reducing conditions (Ishii et al., 2011). Nevertheless, the TPM of *Rhodocyclales nosZ* did not increase in microcosms without N_2_O amendment (Figs. 1d and 2). Overall, it remains unclear why *Rhodocyclales nosZ* is special.

From the viewpoint of social needs for eliminating N_2_O gas, the present study may suggest that *Betaproteobacteria* holding clade II *nosZ* are potential targets of bioaugmentation and biostimulation. In fact, a previous study showed that clade II *nosZ* of *Rhodocyclales* might work as N_2_O sinks in wastewater plants (Kim et al., 2022; Maeda et al., 2024). Although betaproteobacterial *nosZ* is not dominant in native soils (Fig. 5b), it could reside or be activated in soils because their 16S rRNA gene sequences are prevalent, if not dominant, in soil metagenomes (Fig. 5c). In addition, some strains of *Rhodocyclales* have plant growth-promoting functions (Sakoda et al., 2019; Fernández-Llamosas et al., 2020) as well as nitrogen fixation activities (Reinhold-Hurek et al., 1993).

A limitation of this study is the major difference between the field environments and microcosms. Although the amount of N_2_O amended is comparable with N_2_O flux from field soils, N_2_O emitted from outfield soils rapidly disperses into the air. An N_2_O concentration of 4.46 mM is unrealistic in field environments and is far higher than the typical *K_m_* (the so-called Michaelis constant) of NosZ (Yoon et al., 2016; Suenaga et al., 2019; Wang et al., 2023). We may consider that the N_2_O consumption observed in this study represents an upper limit of NosZ produced in microcosms.

In conclusion, although the presence of clade II *nosZ* is not unexpected in soils, the exceptional dominance of *nosZ* transcripts derived from *Rhodocyclales* in response to N_2_O amendment provides insights into the microbial diversity of N_2_O reduction in paddy soils. Our results suggest that *Rhodocyclales* plays a pivotal role in mitigating N_2_O emissions, which has important implications for sustainable agricultural practices and greenhouse gas management strategies, such as biostimulation and bioaugmentation. *In vitro* assessments revealed that *Rhodocyclales* members tends to exhibit higher N_2_O reduction activity among the cultured strains (Hiis et al., 2024), suggesting that stimulating *Rhodocyclales* in soils may enhance soil N_2_O uptake. If strains with superior persistence and colonization capabilities in soil environments can be identified, introducing *Rhodocyclales* to agricultural soils prone to N_2_O emissions could help reduce N_2_O release, as seen in a recent example (Hiis et al., 2024). Future studies should explore the ecological factors driving the selective activation of *Rhodocyclales* in soils and examine how this activation may contribute to the overall functioning of paddy soil microbiomes.

## Data Availability

The metagenomic and metatranscriptomic data obtained in this study have been deposited in DDBJ DRA under the accession numbers DRA019863, DRA019864, and DRA019865. See Table S1 for details. Source codes for the bioinformatic analyses are available in FigShare (https://figshare.com/s/5eb9d7d02e35ea07102a).

## Acknowledgements

This work was financially supported by JPNP18016 commissioned by the New Energy and Industrial Technology Development Organization (NEDO) and JST-Mirai Program grant JPMJMI20E5. We thank Momoko Hamatani, Emiko Kobayashi, Kyohei Kuroda, and Aika Sawaguchi (National Institute of Advanced Science and Technology [AIST]) for supporting the study and Hirotomo Ohba (Niigata Agricultural Research Institute) and anonymous farmers for providing the soil samples. Computations were partially performed on the Bioresource Analysis Platform supercomputer implemented in the AIST Hokkaido and SHIROKANE supercomputers at the Human Genome Center, The Institute of Medical Science, The University of Tokyo.

## Ethical Statement

This study involves no experiments or analyses requiring ethical approval.

## Author Contributions

KM: methodology, software, validation, data curation, formal analysis, investigation, resources, writing: original draft, writing: review and editing, visualization. YM: conceptualization, methodology, validation, investigation, resources, writing: review and editing, funding acquisition. KS: conceptualization, resources, writing: review and editing, supervision, funding acquisition. HI: conceptualization, methodology, validation, investigation, resources, writing: review and editing, supervision, funding acquisition.

