## Supplemental Text, Tables S1 and S2, Figures S1 and S2 for "Betaproteobacterial clade II *nosZ* activated under high N_2_O concentrations in paddy soil microcosms"

**Supplementary Text 1: Full details for phylogenetic and taxonomic annotations of NosZ**

To construct a reference database for phylogenetic placement, we first identified NosZ sequences encoded on GTDB genomes. We linked all genomes (n = 113,104) in GTDB Release 220 (Parks et al., 2022) and their NCBI Taxonomy IDs (Federhen et al., 2012), and eliminated those lacking taxonomic annotations at phylum, class, or order levels. For the remaining 74,660 genomes with valid taxonomic annotations, we identified protein coding sequences (CDSs) using prodigal v2.6.3 (Hyatt et al., 2010) with default parameters. We aligned the CDSs with KOfam profile of K00376 (i.e., the K number representing NosZ), using KofamScan with default parameters (Aramaki et al., 2020). The identified NosZ-encoding CDSs were further classified into two distinct groups, namely clade I NosZ and clade II NosZ. Here we used HMM profiles for clade I NosZ and clade II NosZ provided by FunGene (Fish et al., 2013). Although the web interface of FunGene is currently out of operation, the HMM profile is available (https://github.com/rdpstaff/fungene_pipeline/blob/master/resources). The NosZ-encoding CDSs were aligned with the two HMM profiles using HMMER 3.4 (Eddy, 2007) with default parameters and classified according to bitscores of the alignments. We obtained 7,046 NosZ sequences, each accompanied by two orthogonal information: NCBI Taxonomy ID and binary grouping of NosZ (either clade I [represented by nosZ/model.hmm in FunGene] or clade II [nosZ_a2/model.hmm]). This set of sequences was used as the reference database for annotating NosZ on metatranscriptomic and metagenomic sequences.

The taxonomic annotation of NosZ sequences was performed in the same manner as Masuda et al. (2024) with modifications in software versions. The NosZ database was dereplicated and fed into MAFFT v7.525 (with "--auto" option) (Katoh et al., 2002) to construct a multiple sequence alignment (MSA). Based on the MSA, an approximate maximum-likelihood tree was obtained using FastTree 2.1.11 (Price et al., 2009) with default parameters. The MSA and the phylogenetic tree were used as backbones in phylogenetic placement. NosZ sequences on metatranscriptomic contigs or metagenomic reads (see main text) were mapped onto the backbone MSA using "--add" command in MAFFT. The mapped MSA was subjected to pplacer v1.1.alpha19-0-g807f6f3 (Matsen et al., 2010) to determine the taxonomy and phylogeny of each query sequence.

**Table S1.** Accession numbers of metatranscriptomic and metagenomic data deposited in DDBJ DRA

| Soil name | Metatranscriptome | Metagenome |
|---|---|---|
| X | Day 0, without $N_2O$: DRR624459 – DRR624462 | DRR624512, |
| | Day 2, without $N_2O$: DRR624463 – DRR624465 | DRR624513, |
| | Day 4, without $N_2O$: DRR624466 – DRR624468 | DRR624514 |
| | Day 7, without $N_2O$: DRR624469 – DRR624471 | |
| | Day 1, +$N_2O$: DRR624472 – DRR624475 | |
| | Day 2, +$N_2O$: DRR624476 – DRR624479 | |
| | Day 3, +$N_2O$: DRR624480 – DRR624483 | |
| | Day 4, +$N_2O$: DRR624484 – DRR624487 | |
| | Day 5, +$N_2O$: DRR624488 – DRR624491 | |
| | Day 6, +$N_2O$: DRR624492 – DRR624495 | |
| | Day 7, +$N_2O$: DRR624496 – DRR624499 | |
| Y1 | DRR624500 – DRR624502 | DRR624515 |
| Y2 | DRR624503 – DRR624505 | DRR624516 |
| Y3 | DRR624506 – DRR624508 | DRR624517 |
| Y4 | DRR624509 – DRR624511 | DRR624518 |

**Table S2.** Statistical summary of metatranscriptomic sequencing and assembly for Soils Y1–Y4.

| Soil | Replication | Number of reads* | Number of contigs (>=500bp) | Number of contigs bearing *nosZ* | Mapped reads (proportion) |
|---|---|---|---|---|---|
| Y1 | 1 | 36,277,178 | 321,311 | 125 | 26,009,147 (71.7%) |
| | 2 | 29,503,388 | | | 20,187,502 (68.4%) |
| | 3 | 33,764,380 | | | 21,334,559 (63.2%) |
| Y2 | 1 | 35,548,854 | 450,725 | 145 | 26,409,843 (74.3%) |
| | 2 | 33,817,066 | | | 21,120,651 (62.5%) |
| | 3 | 43,073,028 | | | 25,245,705 (58.6%) |
| Y3 | 1 | 40,255,788 | 328,253 | 138 | 29,953,935 (74.4%) |
| | 2 | 32,922,968 | | | 25,098,778 (76.2%) |
| | 3 | 41,489,496 | | | 22,580,287 (54.4%) |
| Y4 | 1 | 29,520,284 | 306,193 | 228 | 20,529,841 (69.5%) |
| | 2 | 30,037,780 | | | 21,314,871 (71.0%) |
| | 3 | 32,597,386 | | | 22,498,141 (69.0%) |

*: Number of reads after quality filtering and removal of rRNA-like reads

**Figure S1.** Examples of operon structures of *nosZ*-bearing contigs. Each square indicates one coding sequence (CDS), and circled letters indicate functional annotations. CDSs encoding *nosZ* are displayed in orange. Only CDSs on the same direction as *nosZ* are displayed.

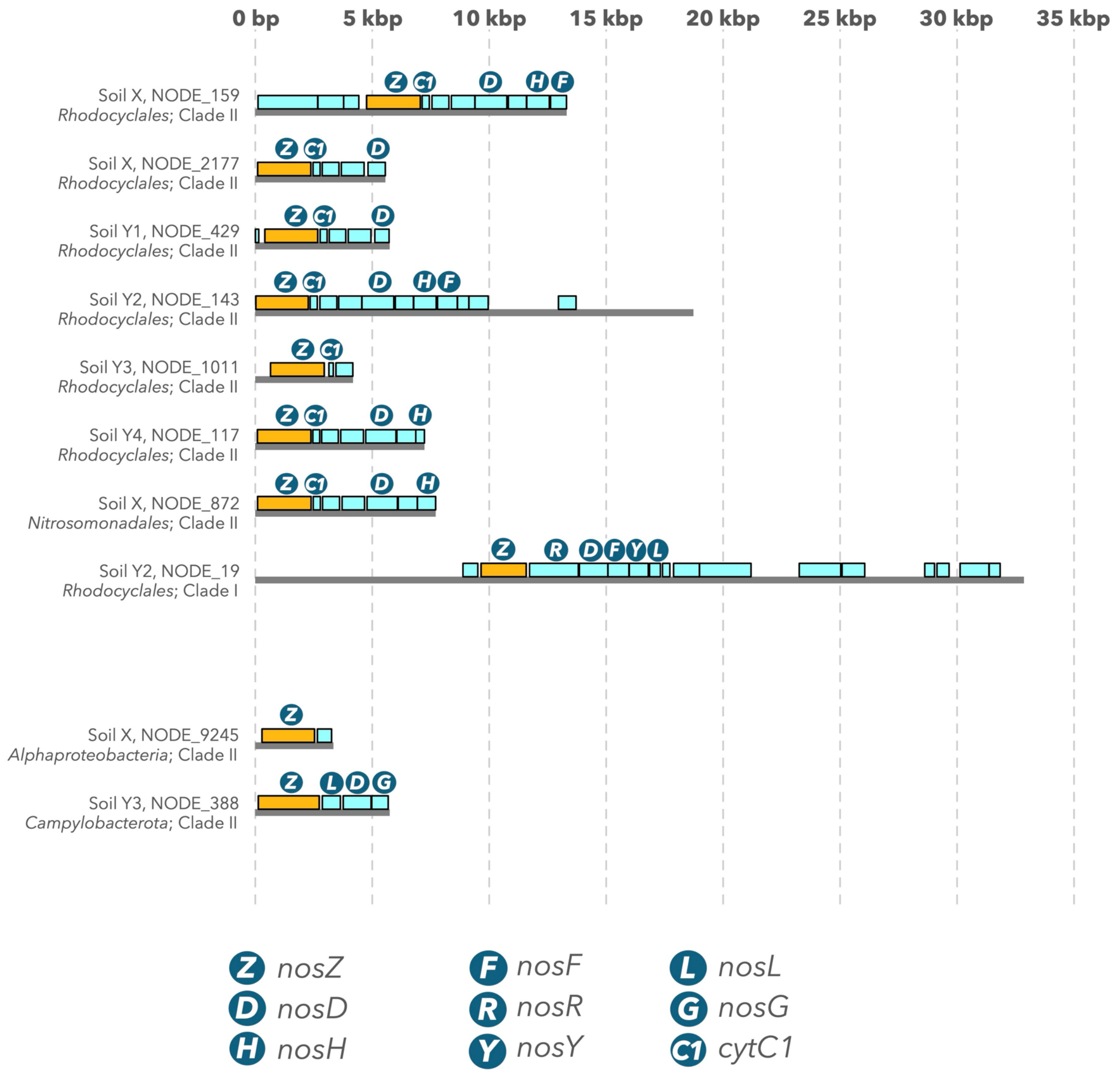

**Figure S2.** Dynamics of transcripts per kilobase million (TPM) of (a) *nosD*, (b) *nosF*, and (c) *nosH* in the metatranscriptomic sequences of Soil X microcosms. Each panel consists of two bar charts, representing samples without and with $N_2O$ amendment. NA denotes not available.

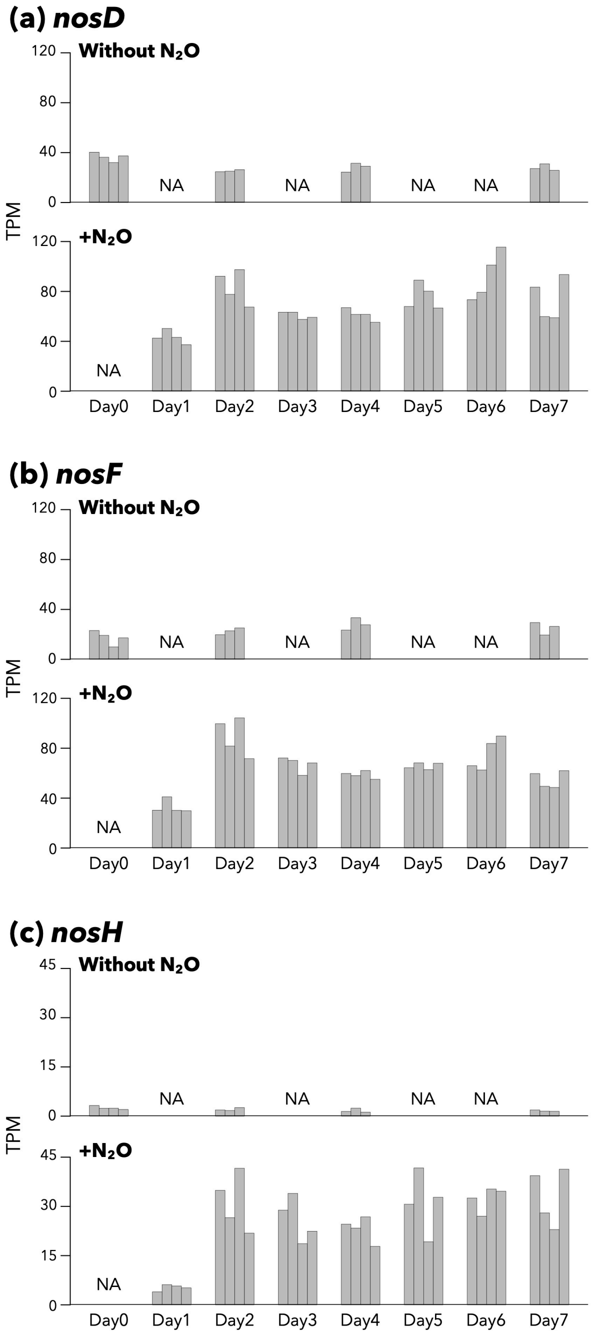
